# Deep vascular imaging in the eye with flow-enhanced ultrasound

**DOI:** 10.1101/2021.06.04.447055

**Authors:** Christian Damsgaard, Henrik Lauridsen

## Abstract

The eye’s retina is one of the most energy-demanding tissues in the body and thus requires high rates of oxygen delivery from a rich blood supply. The capillary lamina of the choroid lines the outer surface of the retina and is the dominating source of oxygen in most vertebrates, but this vascular bed is challenging to image with traditional optical techniques due to its position behind the highly light-absorbing retina. Here we describe a high-frequency ultrasound technique with flow-enhancement to image deep vascular beds (0.5 – 3 cm) of the eye with a high spatiotemporal resolution. This non-invasive method works well in species with nucleated red blood cells (non-mammalian and fetal animal models), and it generates non-invasive three-dimensional angiographies without the use of contrast agents that is independent of blood flow angles and with a higher sensitivity than Doppler based ultrasound imaging techniques.

## INTRODUCTION

The high metabolism on the vertebrate retina imposes an intrinsic tradeoff between two contrasting needs; high blood flow rates and a light path devoid of blood vessels. To avoid visual disturbance of perfusing red blood cells, the retina of all vertebrates receives oxygen and nutrients via a sheet of capillaries *behind* the photoreceptors, the choriocapillaris^1–3^. However, this single source of nutrients and oxygen imposes a diffusion limitation to the thickness of the retina^4,5^, so many visually active species possess a variety of elaborate vascular networks to provide additional blood supply to this metabolically active organ^6^. These vascular beds include blood vessels perfusing the internal retinal layers in mammals and some fishes^4,7–10^, blood vessels on the inner (light-facing) side of the retina found in many fishes, reptiles, and birds^4,11–13^, and countercurrent vascular arrangements of the fish choroid, the choroid *rete mirabile*, that allows for the generation of super-atmospheric oxygen partial pressures^14–20^. Despite that these additional non-choroidal paths for retinal nutrient supply play an essential role in fueling the metabolic requirements of superior vision^4^, the three-dimensional anatomy of these vascular structures is poorly understood, limiting our understanding of the morphological evolution of the vertebrate eye.

Traditionally, retinal blood supply has been studied using optical techniques, such as fundus ophthalmoscopy. This category of techniques provides high-throughput non-destructive information on non-choroidal blood vessel anatomy in high-resolution^21^ and is therefore readily used in clinical diagnosis of abnormalities in retinal vessel structure^22^. However, the photoreceptor layer absorbs the transmitted light and limits the depth of view in these optical techniques, providing reduced information on choroidal structure and function without the use of contrast agent^21^. Similar depth limitations are experienced in optical coherence tomography (OCT), which generates high-resolution fundus angiographies using light waves at the technical expense of depth penetration^23^. Magnetic resonance imaging overcomes the optical limitations of ophthalmoscopy and OCT and can map vascular layers in the retina, albeit at a low resolution^24^. Histology and microcomputed tomography (μCT) maintain the high-resolution of the optical techniques and provide information on whole-eye vascular morphology^4^, but both techniques require ocular sampling and are therefore not possible in the clinic or in rare or endangered species. To overcome some of the limitations of established retinal imaging techniques, we here present an ultrasound protocol on anesthetized animals, where blood movement is mapped *in silico* on a series of equally-spaced two-dimensional ultrasound scans spanning a whole eye by applying a comparable technique as described previously for embryonic and cardiovascular imaging^25–27^. This approach allows for the generation of non-invasive three-dimensional deep ocular angiographies without using a contrast agent and opens up new avenues for mapping blood flow distribution within the eye across species.

## PROTOCOL

### 1. Anesthesia and ultrasound medium

1.1. Anesthetize research animal

**Note:** Type and dose of appropriate anesthesia are highly species-dependent. In general, immersion-based anesthetics such as MS-222 (ethyl 3-aminobenzoate methanesulfonic acid), benzocaine (ethyl 4-aminobenzoate), and propofol (2,6-diisopropyl phenol) are useful in fish and amphibians which readily absorbs the anesthetic over gills or skin. A range of dissolved compounds that can be administered intravenously, intramuscularly, intraperitoneally is available for amniotes, as are gas-based anesthetics. In our experience, alfaxalon administered intramuscularly is useful in reptiles and isoflurane administered as gas is useful in birds. We point to ^28–30^ for a full overview of available anesthetics across species.

1.2. Test reflexes in the animal.

**Note:** The flow-enhanced ultrasound procedure is sensitive to motion noise. Thus the animal must be completely motionless during the procedure. However, too deep anesthesia can alter blood flow patterns, so it is advisable to conduct a dose titration in the start-up phase of an experiment where blood flow in the eye is observed aided by simple B-mode ultrasound as anesthesia dosage is increased in steps. An optimal level of anesthesia is obtained when the animal is motionless (except respiration) with normal ocular blood flow patterns.

1.3. If the type/dose of anesthetic is not permissive for respiratory movements, then ensure adequate ventilation of the animal, *e.g.*, using an air pump to oxygenate the water for aquatic species or a ventilator for air-breathing species.

1.4. Position the animal in a posture that allows direct access from above to the eye.

**Note:** Depending on species, this can be in either a supine or lateral position. It can be useful to construct a simple holding device using a small piece of non-reactive metal (*e.g.*, stainless steel) and loose rubber bands (see Fig. 1).

**Figure 1.**
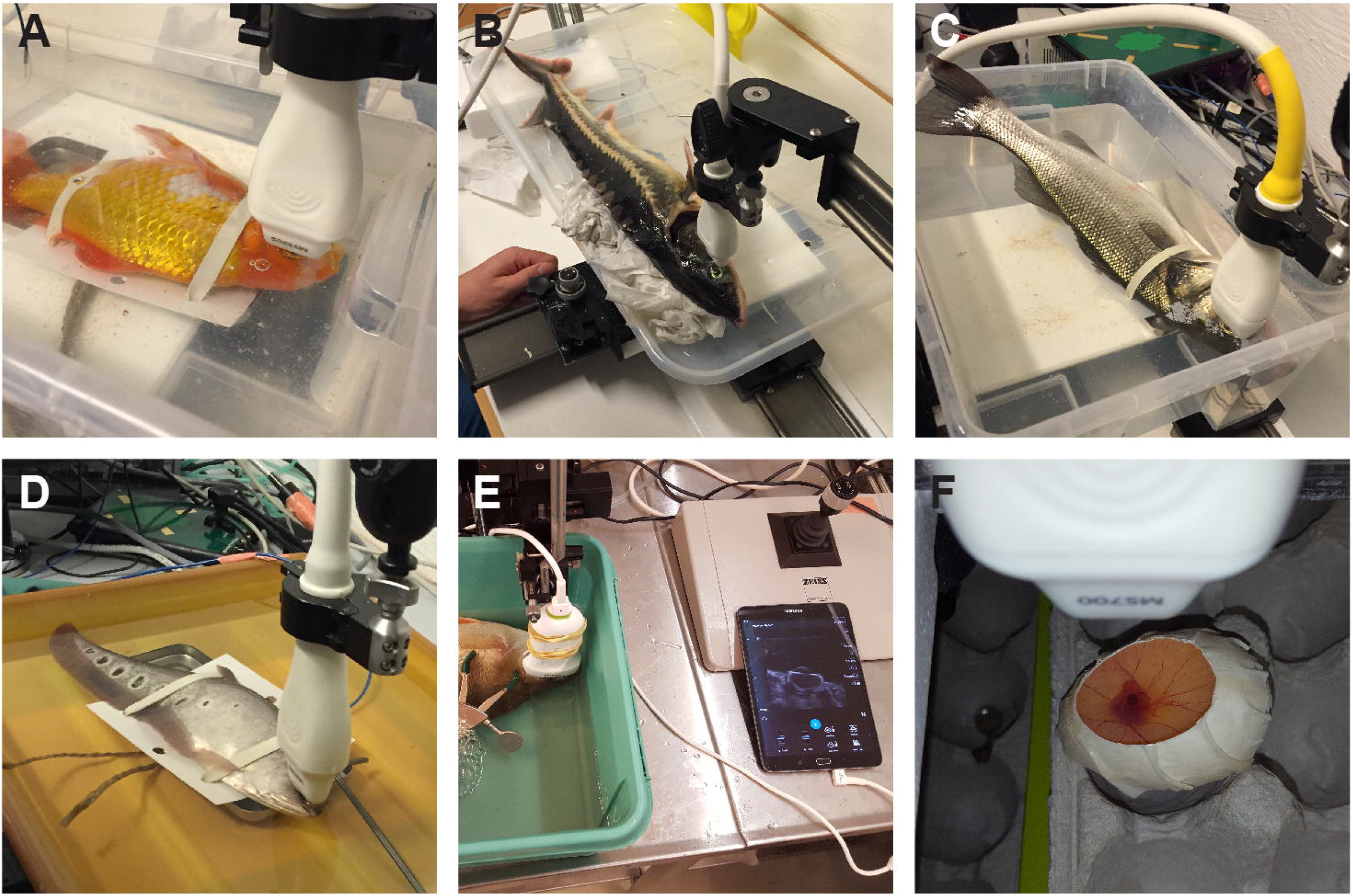
Examples of the variety of species suitable for flow-enhanced ultrasound imaging of ocular vasculature. **A**, goldfish (*Carassius auratus*). **B**, Siberian sturgeon (*Acipenser baerii*). **C**, European seabass (*Dicentrarchus labrax*). **D**, clown featherback (*Chitala ornata*). **E**, Crucian carp (*Carassius carassius*). **F**, embryonic domestic chicken (*Gallus gallus domesticus*). It can be useful to construct a simple holding device using a non-reactive metal weight and loose rubber bands (**A**, **C**, **D**). Both large, immobile lab-based ultrasound imaging systems can be used for the procedure (**A – D**, **F**) as well as small field operative systems (**E**). When imaging small and highly temperature-sensitive species that cannot be retained in a temperature-controlled water bath like embryonic birds, imaging can be performed while the sample is inside the incubator (**F**).

1.5. Place appropriate ultrasound medium on the eye of the animal. If scaled eyelids (ultrasound impermeable) cover the eye, then these should be displaced gently with a cotton swab.

**Note:** For aquatic species, the best ultrasound medium is clean tank water in which the animal usually lives. For terrestrial species, a generous amount of ultrasound gel ensures free movements and imaging of the ultrasound transducer across the entire surface of the eye.

### 2D and 3D ocular ultrasound image acquisition

2.1. Position ultrasound transducer medial to the eye in either a dorsal/ventral or rostral/caudal orientation depending on desired image orientation.

2.2. In B-mode with maximum depth of field, image the medial and deepest portion of the eye and make sure that all structures of interest are visible in the image field.

2.3.Slowly translate the transducer to each side while inspecting the real-time images. Make sure all structures of interest are visible in the image field; if not, switch to a transducer with a lower frequency and larger depth of field.

**Note:** In our experience, the following center frequencies allow for the following maximum depth of field: 21 MHz: 3 cm, 40 MHz: 1.5 cm, 50 MHz: 1 cm. However, these maximum depth of field values can be markedly lower if the eye contains calcified or other ultrasound impermeable structures.

2.4. Adjust image depth, depth offset (distance from the top of the image to structure of interest), image width, as well as number and position of focal zones to cover the desired region of interest in all three spatial dimensions.

**Note:** These image parameter settings usually affect the range of possible temporal resolutions of the ultrasound acquisition.

2.5. Set frame rate in the range of 50 – 120 frames s^−1^.

**Note:** The temporal resolution must be adequate to display large pixel intensity variability in imaged blood vessels, *i.e.*, the temporal resolution must not be too high. On the other hand, to complete a full 3D recording of the eye in a reasonable time, temporal resolution cannot be too low. In our experience, a temporal resolution ranging from 50 – 120 frames s^−1^ is usually adequate for the flow-enhanced procedure in most species. On some ultrasound systems, this desired temporal resolution can be obtained by switching between the “general imaging” (high spatial/low temporal resolution) and “cardiology” (low spatial/high temporal resolution) modes.

2.6. Adjust 2D gain to a level (~5 dB), so anatomical structures are only just visible in the B-mode acquisition to increase the signal-to-noise ratio in the subsequent flow-enhanced reconstruction.

2.7.To acquire a 2D flow-enhanced image at a single slice position, translate the transducer to this position and continue at step 3.1.

2.8. To acquire a 3D recording of an entire region of interest, *e.g.*, the retina, translate the transducer to one extreme of the region of interest.

**Note:** To determine the exact position of the extreme end of the region of interest, it may be necessary to increase the 2D gain briefly. After correct transducer placement has been completed, the 2D gain must be lowered before recording to ensure maximal signal-to-noise ratio in the subsequent flow-enhanced reconstruction.

2.9. For each step (slice) in the 3D recording, acquire ≥100 frames (optimally ≥1000 frames).

2.10.Using a micromanipulator or build-in transducer motor, translate the transducer across the entire region of interest in steps of, *e.g.*, 25 or 50 μm (remember to note the step size) and repeat the ≥100 frames acquisition for each step.

### 3. Flow-enhanced image reconstruction

3.1. Export recordings into digital imaging and communications in medicine (DICOM) file format (little endian).

3.2. To produce a single flow-enhanced image based on a ≥100 frames (*T*) cine recording, calculate the standard deviation on pixel level (*STD(x,y)*) using the formula:

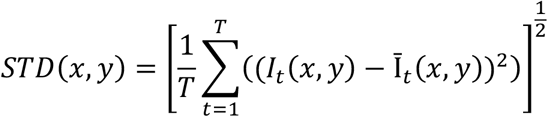

where *I_t_(x,y)* is the intensity of the pixel at the *(x,y)* pixel coordinate at time *t*, and *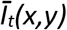* is the arithmetic mean value of *I* over time.

3.3. Repeat step 3.2 for each slice in the 3D recording.

**Note:** To automate the STD-calculation and image reconstruction process for multiple slices in a 3D recording, this operation can be conducted in batch mode using, *e.g.*, ImageJ and the supplementary macro script (Supplementary file 1).

3.4. Combine all reconstructed slices into one image stack.

3.5. Specify slice thickness from the step size used during acquisition.

3.6. Save image stack as a 3D TIF file.

**Note:** Flow-weighted three-dimensional recordings of ocular blood vessels can subsequently be used to create volume renderings and build digital and physical anatomical models of vascular structures of the eye. These image processing options are outside the scope of this protocol, and we instead point to^31–33^.

## REPRESENTATIVE RESULTS

The flow-enhanced ultrasound technique to image vascular beds of the eye can be applied in a range of species, and we have currently used it in 46 different vertebrate species (Fig. 1, Table 1). The presence of nucleated red blood cells in non-adult-mammalian vertebrates provides positive contrast of flowing blood compared to static tissue in cine recordings (Supplementary file 2). However, when analyzed on a frame-by-frame basis, the clear distinction between blood and surrounding tissue is less obvious (Fig. 2A). The blood flow enhancement procedure described in this protocol essentially compiles a multi-time point recording in 2D space (a slice made of *T* frames) into a single image in which the inherent signal value fluctuations in pixels positioned in flowing blood scores a higher standard deviation than surrounding static tissue, hence producing positive contrast (Fig. 2B). To perceivably enhance the blood vessel contrast, Look Up Tables can be used to produce pseudocolor images (Fig. 2C). In 3D acquisitions, multiple parallel slices with known spacing can be combined into 3D image data (Supplementary file 3 and 4) that can be used for three-dimensional volume rendering (Fig. 2D) and anatomical modeling (Fig. 2E and Supplementary file 5). Doppler-based ultrasound imaging also provides the option to specifically image blood flow, however with less sensitivity than the described method (compare Fig. 2G with Fig. 2H and 2I), and importantly not if blood flow orientation is directly or close to perpendicular to the direction of the sound wave. The flow-enhanced procedure described in this protocol is independent of the orientation of blood flow both in-plane and out-of-plane.

**Figure 2.**
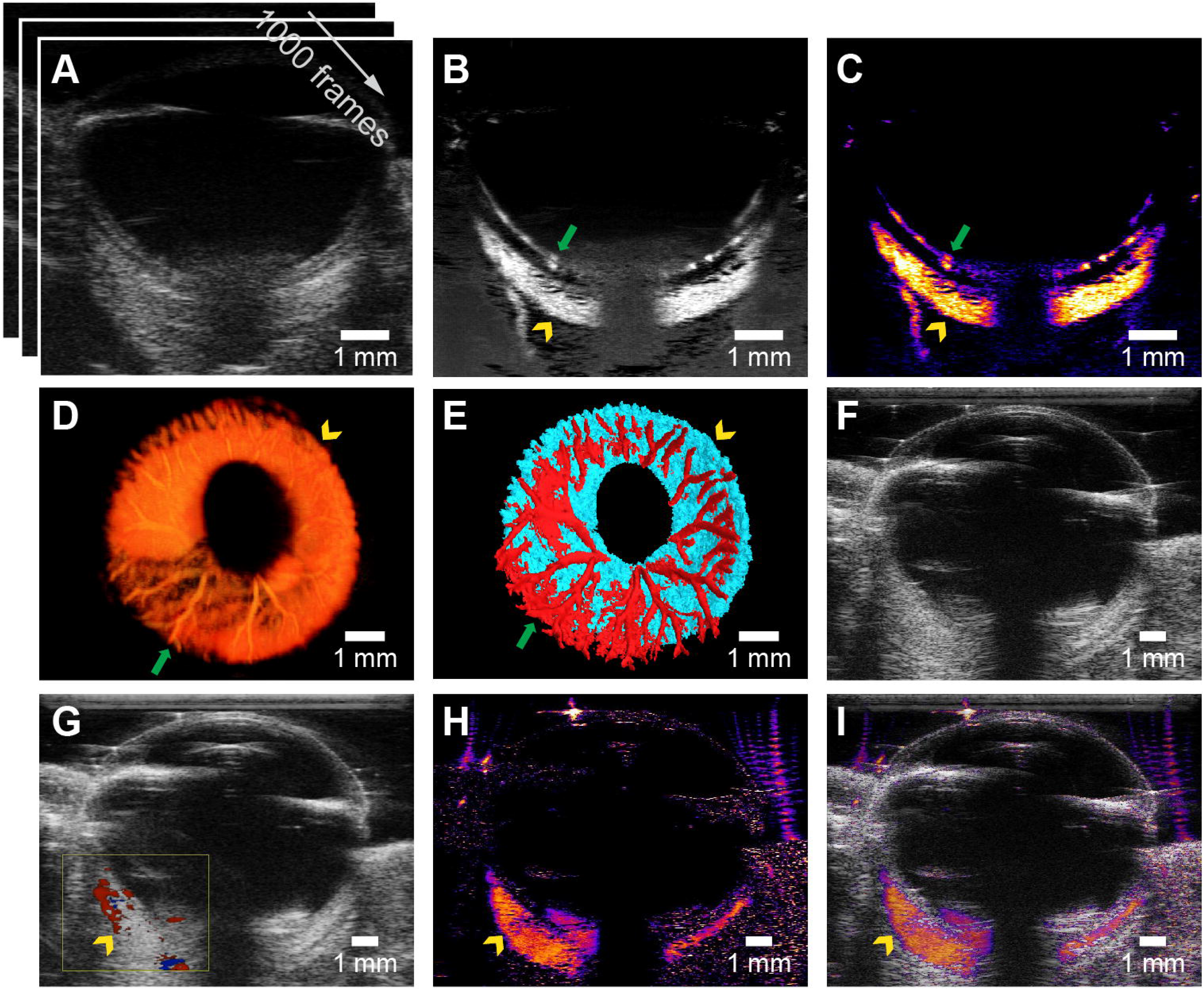
Effect of flow-enhancement. **A**, Examples of raw B-mode ultrasonographic images of the eye of a goldfish in a 1000 frame cine recording. Whereas blood flow can be observed in the cine recording (supplementary material 2) it is difficult to see in static frames. **B**, flow-enhanced grayscale image (same slice as in **A**). Both retinal and post-retinal vascular beds are enhanced. **C**, pseudo-colored version of the image in **B** with ImageJ Fire Look Up Table. **D**, volume-rendered display of blood flow in the eye of the same goldfish as in **A-C**, based on 3D acquisition. **E**, two-segment (retinal and post-retinal vessels) anatomical model of eye in **A-D** (for interactive model see supplementary material 5). **F-I**, raw B-mode ultrasonographic image of the eye of another goldfish (**F**) comparing color Doppler based flow imaging (**G**) to the flow-enhanced methods described in this protocol (**H-I**, note **I** is an overlay of **H** on **F**). Green arrows indicate retinal vessels, yellow arrowheads indicate the choroid *rete mirabile*.

The flow-enhanced ultrasound procedure allows for blood flow imaging in a range of species with nucleated red blood cells (Fig. 3A - 3D). Deep ocular vascular beds such as the choroid *rete mirabile* in some fish can be imaged if present in the species (yellow arrowhead in Figs. 2, 3B,4). The method is limited by the absence of nucleated red blood cells in adult mammals in which the flow enhancement procedure produces some degree of blood flow contrast but not as distinct as in species with nucleated red blood cells (Fig. 3E and 3F).

**Figure 3.**
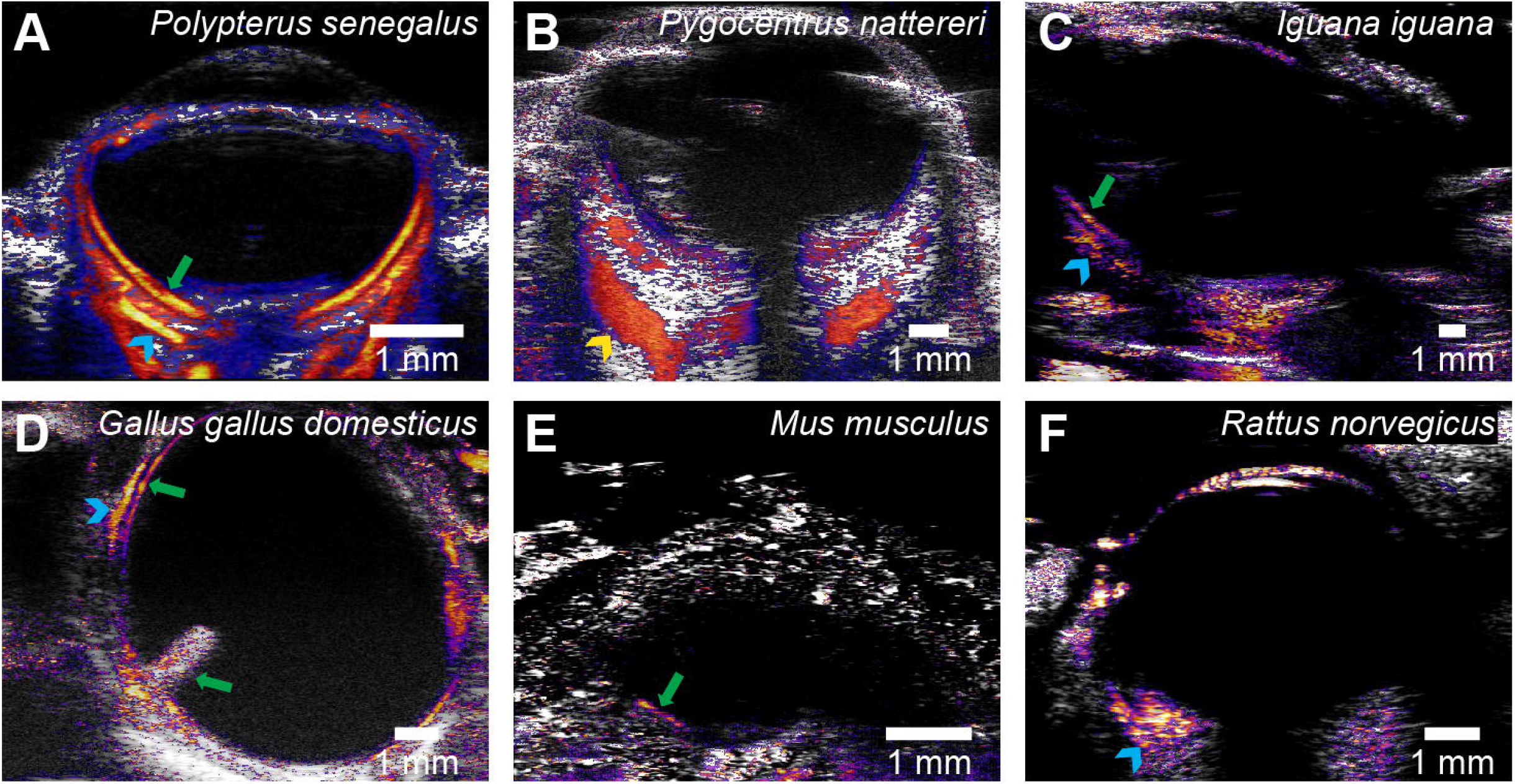
Representative examples of flow-enhanced ocular ultrasound images in a variety of vertebrate species. **A**, Senegal bichir (*Polypterus senegalus*). **B**, red-bellied piranha (*Pygocentrus nattereri*). **C**, green iguana (*Iguana iguana*). **D**, embryonic (day 18) domestic chicken (*Gallus gallus domesticus*). **E**, House mouse (*Mus musculus*). **F**, brown rat (*Rattus norvegicus*). In species with nucleated red blood cells, the flow-enhancement procedure yields useful images of ocular blood flow (**A-D**), whereas in adult mammals (enucleated red blood cells), it produces only limited contrast between flowing blood and surrounding tissue (**E-F**). Green arrows indicate retinal vessels, blue arrowheads indicate post-retinal vessels such as the choriocapillaris, yellow arrowheads indicate choroid *rete mirabile*. In the late embryonic domestic chicken, blood flow in the pecten oculi can be observed (lower green arrow in **F**).

Flow-enhanced ultrasound is sensitive to motion noise, and *e.g.*, respiratory movements can course image blurring and artifacts such as tissue border enhancement (Fig. 4A – 4C, Supplementary file 6). Prospective or retrospective gating can be used to adjust for motion noise (Fig. 4D and 4E).

**Figure 4.**
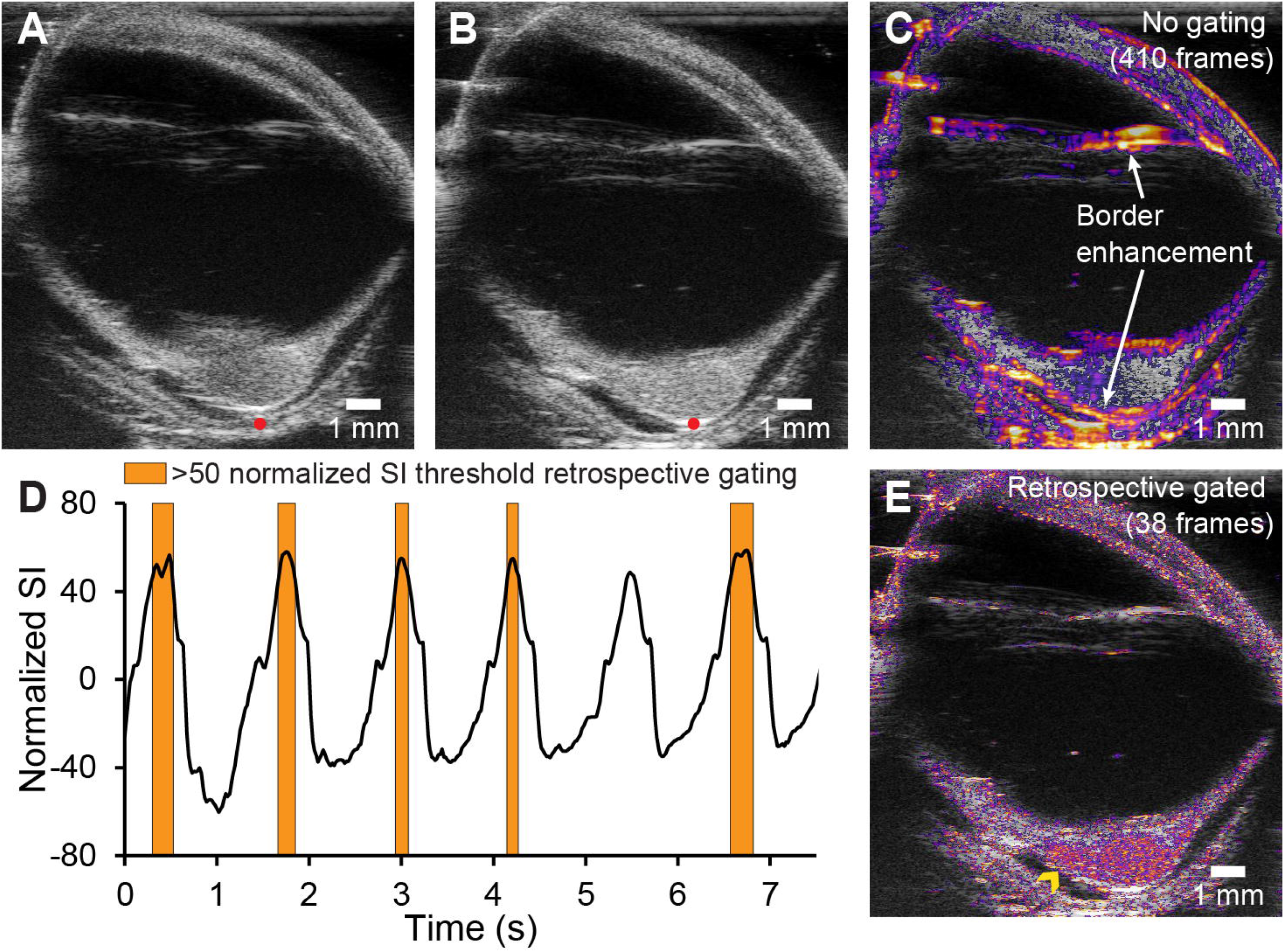
Respiratory movements induce motion noise that can be alleviated by retrospective gating. **A-B**, Example of respiratory movements in the eye of a European plaice (*Pleuronectes platessa*). Red dot is at the same image coordinate in **A** (slice 54/410) and **B** (slice 92/410), but it can be observed that the eye has shifted position (see also cine recording in supplementary material 6). **C**, attempt to perform flow-enhancement operation on the full 410 frames recording fails due to motion noise. Tissue borders are artificially enhanced due to movements. **D**, retrospective gating operation based on normalized signal intensity (SI) at the red dot in **A-B**. Only frames with normalized SI > 50 (in total 38 frames), *i.e.*, indicating that the eye is at the same position as in **B,** are included for the flow-enhancement procedure. **E**, resulting image of retrospectively gated flow-enhancement procedure. Compare with **C**. In the gated image, artificial border enhancement is avoided, and blood flow in the choroid *rete mirabile* (yellow arrowhead) can be observed.

**Table 1.** List of species that the flow-enhanced ultrasound technique to image ocular blood flow has been used on. The applicability of the method is based on the ability to produce contrast rich representation of vascular beds compared to static background.

## DISCUSSION

Vascular imaging using flow-enhanced ultrasound provides a new method for non-invasive imaging of the vasculature of the eye that offers several advantages over present techniques but has its intrinsic limitations. The primary advantage of flow-enhanced ultrasound is the ability to generate ocular angiographies with a depth of field that exceeds the photoreceptor layer, which limits the depth of field in optical techniques. In ultrasound imaging, spatial resolution and depth of field are ultimately determined by the ultrasound transducer frequency, where higher frequencies increase the spatial resolution but at the expense of a shallower depth of field, thus the choice of transducer frequency introduces a tradeoff between image depth and spatial resolution. In our experience, optimal retinal ultrasound imaging is achieved using high-frequency ultrasound transducers (≥50 MHz) in small eyes with image depths of <1 cm and lower frequency transducers (20-40 MHz) in larger eyes with image depths of 1.5 – 3.0 cm. For a 3D ultrasound scan, the resolution of the additional slice-dimension is set by the step size between scans in the stack of 2D ultrasound scans. In our experience, it is difficult to conduct a 3D scan with a step size smaller than 20 μm.

Flow-enhanced 2D ultrasound has a high temporal resolution. Ideally, ≥1000 frames per image are required for flow-enhanced vascular imaging, so at least 8 seconds are required per image scan. The temporal resolution is significantly reduced when performing 3D flow-enhanced ultrasound, where the scanning time increases with the number of images in the 3D stack of scans. Given the high temporal resolution, the flow-enhanced 2D ultrasound workflow shows strong potential as a method for identifying temporal changes in relative blood flow velocities and blood flow distribution during experimental manipulation. Thus, future studies can use the workflow to identify how altered environmental temperatures (*e.g.*, temperature, *p*O_2_, *p*CO_2_) or pharmacological administration affect blood flow in the eye and other organs.

The ultrasound workflow relies on the positive contrast of nucleated red blood cells from most non-mammalian vertebrates. Thus, the enucleated red blood cells of mammals and some salamander species^34^ provide too little contrast to effectively enhance blood flow using the present workflow (Fig. 3EF). In traditional ultrasound workflows, vascular injection of microbubbles provides high enough contrast to identify the vasculature in mammals^35^. However, our initial attempts to generate flow-enhanced angiographies in rodents using vascular microbubble injection failed, so future implementation of flow-enhanced ultrasound mapping of mammalian vasculatures replies on the optimization microbubble dosages and ultrasound settings^35^.

Flow-enhanced ultrasound depends on sequential recordings in the same position of the eye, so the technique is not possible in awake animals, where minor random movements may offset the image and undermine flow-enhancement calculations. Thus, the present method must be performed under proper anesthesia for immobilization to enhance image quality by reducing random movements. However, regular movements of the eye that occur during regular respiratory movements can be offset by prospectively or retrospectively gating to the ventilation pattern of the animal, so only scan recording from the same time interval within the ventilation cycle is used in the data analysis. While the retrospective gating approach to offset ventilatory movements of the image significantly improves the image stability, it pronouncedly reduces the number of frames included in calculating standard deviation of signal intensity leading to a decrease in signal-to-noise ratio (compare Fig. 4E to Fig. 2C and 2I). This effect is alleviated using prospective gating at the ultrasound scanner in which image data is only acquired when the animal is in the desired phase of respiration. However, this causes a marked increase in acquisition time if a desired number of frames ≥1000 must be acquired.

We see multiple applications in zoological and veterinarian research for the flow-enhanced ultrasound workflow to map the physiology and anatomy of the eye’s vasculature. The vasculature of ray-finned fishes, mammals, and birds are relatively well-described^1,3,4,8,9,12,15,36^, but this is not the case for non-bony fishes (jaw-less vertebrates and chondrichthyans), amphibians, and reptiles, that represent their respective earlier diverging sister groups. Implementing flow-enhanced ultrasound on these poorly understood animal groups and integrating these data with knowledge on the more well-studies groups will provide fundamental insight into the evolution of the vasculature of the vertebrate eye. Because the eye’s vasculature is similar in closely related species^4^, such detailed information on the ocular vasculature in a broad range of species will provide a point-of-reference for veterinarians to identify malformations in the eye’s vasculature due to developmental defects, diseases, or physical injuries. Furthermore, the ability to acquire 2D blood flow information with a high spatiotemporal resolution provides the means for quantifying pharmacokinetic effects on blood flow distribution and velocities in deep vascular beds, with vast applications in drug development and testing. Future studies on this technique should focus on identifying injectable compounds that enhance the contrast of blood in species with enucleated red blood cells, which will expand the applicability of this technique to mammals with vast applications in biomedical research and clinical diagnostics of vascular dysfunction in the eye and other deep vascular beds.

## SUPPLEMENTARY FILES

**Supplementary file 1.** Macro script to automate flow-enhancement calculations. Script is written in IJ1 Macro language and can be executed both using the ImageJ macro function (for single slice recording) or the ImageJ Batch Process (for multiple slice 3D recording).

**Supplementary file 2.** Raw B-mode cine recording on the eye of a goldfish (*Carassius auratus*). Blood flow can be observed as the video is playing, but not on a single frame as in Fig. 2A.

**Supplementary file 3.** Slice video through the eye of a goldfish (*Carassius auratus*) of blood flow-enhanced sections.

**Supplementary file 4.** Three-dimensional TIF file of flow-enhanced eye of goldfish (*Carassius auratus*). Images have been binned by 3 × 3 × 3 to minimize file size (27-fold reduction in spatial resolution and file size).

**Supplementary file 5.** Interactive 3D model of pre- and post-retinal vessels in the eye of a goldfish (*Carassius auratus*).

**Supplementary file 6.** Raw B-mode cine recording on the eye of a European plaice (*Pleuronectes platessa*). Note respiratory movements.

## ACKNOWLEDGMENTS

This work has received funding from the Carlsberg Foundation (CF17-0778; CF18-0658), the Lundbeck Foundation (R324-2019-1470; R346-2020-1210), the Velux Foundations (00022458), The A.P. Møller Foundation for the Advancement of Medical Science, the European Union’s Horizon 2020 research and innovation program under the Marie Skłodowska-Curie grant agreement (No. 754513), and The Aarhus University Research Foundation.

## DISCLOSURES

The authors declare that no completing interests exists.

## Notes

### Competing Interest Statement

The authors have declared no competing interest.

## REFERENCES

1 Yu, C. Q., Schwab, I. R. & Dubielzig, R. R. Feeding the vertebrate retina from the Cambrian to the Tertiary. Journal of Zoology. 278(4), 259–269, (2009).

2 Yu, D. Y. & Cringle, S. J. Oxygen distribution and consumption within the retina in vascularised and avascular retinas and in animal models of retinal disease. Progress in Retinal and Eye Research. 20(2), 175–208, (2001).

3 Country, M. W. Retinal metabolism: A comparative look at energetics in the retina. Brain Research. 167250–57, (2017).

4 Damsgaard, C. et al. Retinal oxygen supply shaped the functional evolution of the vertebrate eye. Elife. 8, (2019).

5 Buttery, R. G., Hinrichsen, C. F. L., Weller, W. L. & Haight, J. R. How thick should a retina be? A comparative study of mammalian species with and without intraretinal vasculature. Vision Research. 31(2), 169–187, (1991).

6 Ames, A., Li, Y., Heher, E. & Kimble, C. Energy metabolism of rabbit retina as related to function: high cost of Na^+^ transport. The Journal of Neuroscience. 12(3), 840–853, (1992).

7 Chase, J. The Evolution of Retinal Vascularization in Mammals: A Comparison of Vascular and Avascular Retinae. Ophthalmology. 89(12), 1518–1525, (1982).

8 Johnson, G. L. Ophthalmoscopic studies on the eyes of mammals. Philosophical Transactions of the Royal Society of London. Series B, Biological Sciences. 254(794), 207–220, (1968).

9 Johnson, G. L. I. Contributions to the comparative anatomy of the mammalian eye, chiefly based on ophthalmoscopic examination. Philosophical Transactions of the Royal Society of London. Series B, Biological Sciences. 194(194-206), 1–82, (1901).

10 Rodriguez-Ramos Fernandez, J. & Dubielzig, R. R. Ocular comparative anatomy of the family Rodentia. Veterinary Ophthalmology. 16(s1), 94–99, (2013).

11 Copeland, D. E. Functional vascularization of the teleost eye. Current Topics in Eye Research. 3219–280, (1980).

12 Meyer, D. B. in The Visual System in Vertebrates. Handbook of Sensory Physiology Vol. 7 (ed F. Crescitelli) (Springer, Berlin, Heidelberg, 1977).

13 Potier, S., Mitkus, M. & Kelber, A. Visual adaptations of diurnal and nocturnal raptors. Semin Cell Dev Biol. 106116–126, (2020).

14 Wittenberg, J. B. & Wittenberg, B. A. Active secretion of oxygen into the eye of fish. Nature. 194106–107, (1962).

15 Damsgaard, C. Physiology and evolution of oxygen secreting mechanism in the fisheye. Comparative Biochemistry and Physiolology. 252A110840, (2021).

16 Damsgaard, C. et al. A novel acidification mechanism for greatly enhanced oxygen supply to the fish retina. Elife. 9, (2020).

17 Wittenberg, J. B. & Haedrich, R. L. The choroid rete mirabile of the fish eye. II. Distribution and relation to the pseudobranch and to the swimbladder rete mirabile. Biological Bulletin. 146(1), 137–156, (1974).

18 Wittenberg, J. B. & Wittenberg, B. A. The choroid rete mirabile of the fish eye. I. Oxygen secretion and structure: comparison with the swimbladder rete mirabile. Biological Bulletin. 146(1), 116–136, (1974).

19 Berenbrink, M. Historical reconstructions of evolving physiological complexity: O2 secretion in the eye and swimbladder of fishes. Journal of Experimental Biology. 210(Pt 9), 1641–1652, (2007).

20 Berenbrink, M., Koldkjaer, P., Kepp, O. & Cossins, A. R. Evolution of oxygen secretion in fishes and the emergence of a complex physiological system. Science. 307(5716), 1752–1757, (2005).

21 Keane, P. A. & Sadda, S. R. Retinal Imaging in the Twenty-First Century: State of the Art and Future Directions. Ophthalmology. 121(12), 2489–2500, (2014).

22 Yung, M., Klufas, M. A. & Sarraf, D. Clinical applications of fundus autofluorescence in retinal disease. International Journal of Retina and Vitreous. 2(1), 12, (2016).

23 Ang, M. et al. Optical coherence tomography angiography: a review of current and future clinical applications. Graefe’s Archive for Clinical and Experimental Ophthalmology. 256(2), 237–245, (2018).

24 Shen, Q. et al. Magnetic resonance imaging of tissue and vascular layers in the cat retina. Journal of Magnetic Resonance Imaging. 23(4), 465–472, (2006).

25 Tan, G. X., Jamil, M., Tee, N. G., Zhong, L. & Yap, C. H. 3D Reconstruction of Chick Embryo Vascular Geometries Using Non-invasive High-Frequency Ultrasound for Computational Fluid Dynamics Studies. Ann Biomed Eng. 43(11), 2780–2793, (2015).

26 Ho, S., Tan, G. X. Y., Foo, T. J., Phan-Thien, N. & Yap, C. H. Organ Dynamics and Fluid Dynamics of the HH25 Chick Embryonic Cardiac Ventricle as Revealed by a Novel 4D High-Frequency Ultrasound Imaging Technique and Computational Flow Simulations. Annals of Biomedical Engineering. 45(10), 2309–2323, (2017).

27 Dittrich, A., Thygesen, M. M. & Lauridsen, H. 2D and 3D Echocardiography in the Axolotl (*Ambystoma Mexicanum*). JoVE. doi:10.3791/57089 (141), e57089, (2018).

28 Clarke, K. W., Trim, C. M. & Trim, C. M. Veterinary Anaesthesia E-Book. (Elsevier Health Sciences, 2013).

29 Flecknell, P. Laboratory Animal Anaesthesia. (Elsevier Science & Technology, 2015).

30 West, G., Heard, D. & Caulkett, N. Zoo Animal and Wildlife Immobilization and Anesthesia. (John Wiley & Sons, Incorporated, 2014).

31 Lauridsen, H., Hansen, K., Nørgård, M. Ø., Wang, T. & Pedersen, M. From tissue to silicon to plastic: three-dimensional printing in comparative anatomy and physiology. Royal Society Open Science. 3(3), 150643, (2016).

32 Lauridsen, H. et al. Inside Out: Modern Imaging Techniques to Reveal Animal Anatomy. PLoS One. 6(3), e17879, (2011).

33 Ruthensteiner, B. & Heß, M. Embedding 3D models of biological specimens in PDF publications. Microscopy Research and Technique. 71(11), 778–786, (2008).

34 Mueller, R. L., Ryan Gregory, T., Gregory, S. M., Hsieh, A. & Boore, J. L. Genome size, cell size, and the evolution of enucleated erythrocytes in attenuate salamanders. Zoology. 111(3), 218–230, (2008).

35 Greis, C. Quantitative evaluation of microvascular blood flow by contrast-enhanced ultrasound (CEUS). Clinical Hemorheology and Microcirculation. 49137–149, (2011).

36 Walls, G. L. The vertebrate eye and its adaptive radiation. (Cranbrook Institute of Science, 1944).

