## Supplementary file 1 for "Deep vascular imaging in the eye with flow-enhanced ultrasound"

/*Reconstructs flow enhanced images of 1000 frame ultrasound acquisition*/

//Crop non-essential parts of the echocardiographic image. Adjust crop-size to actual acquisition.
makeRectangle(287, 107, 565, 511);
run("Crop");

//Covert from 8-bit RGB to 8-bit grayscale and then to 32-bit.
run("8-bit");
run("32-bit");

//Measure pixel-wise standard deviation across all frames. In run("Divide..) command specify the number of frames in acquisition. 

rename("X");
run("Z Project...", "projection=[Average Intensity]");
imageCalculator("Subtract create stack", "X","AVG_X");
selectWindow("Result of X");
run("Square", "stack");
run("Z Project...", "projection=[Sum Slices]");
run("Divide...", "value=1000");
run("Square Root");
selectWindow("Result of X");
close();
selectWindow("AVG_X");
close();
selectWindow("X");
close();
selectWindow("SUM_Result of X");
