## Supplementary figures and images for "Deep vascular imaging in the eye with flow-enhanced ultrasound"

### Supplementary file 5

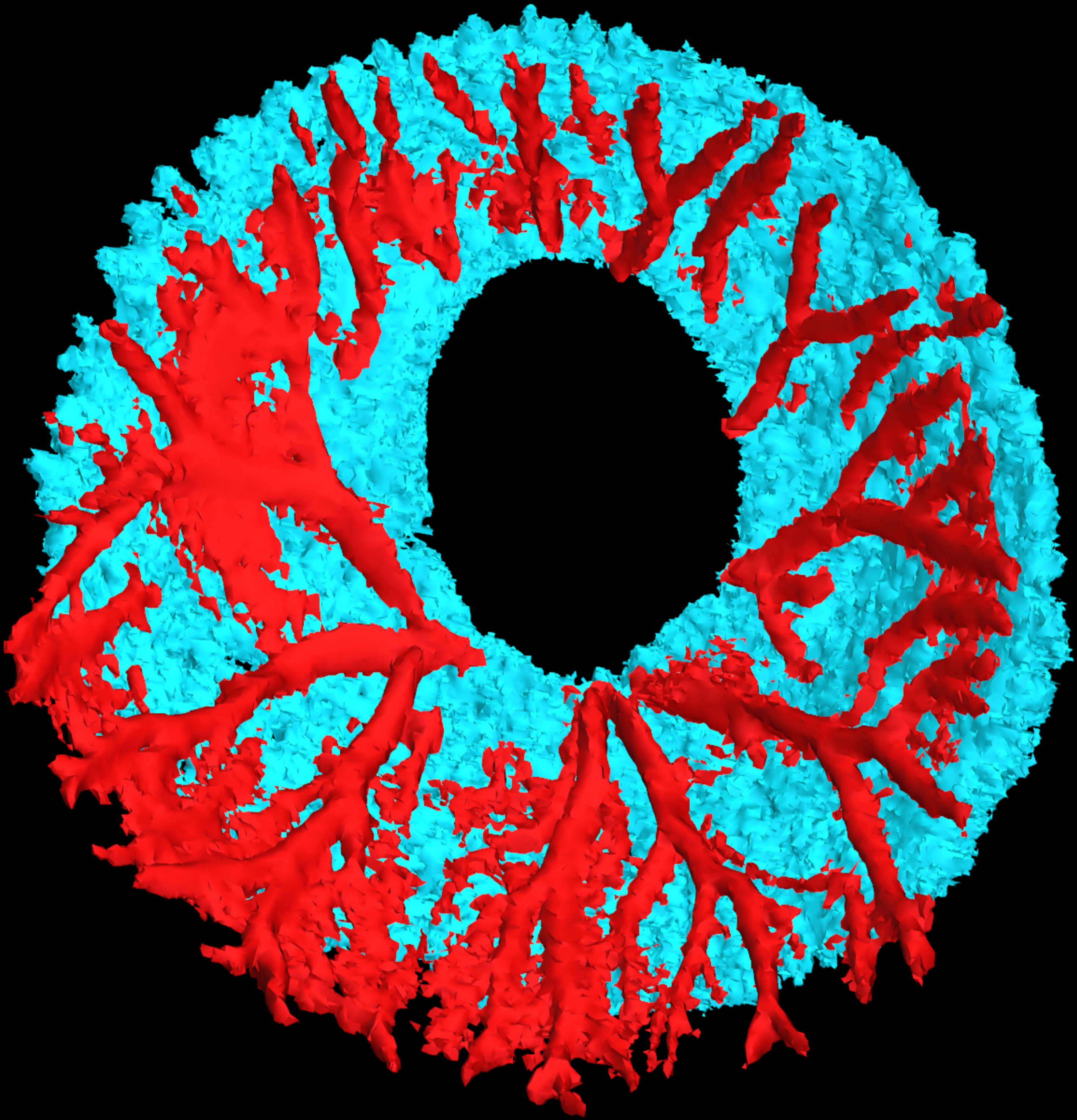
